# A Novel function for Cactus/IκB inhibitor to promote Toll signals in the *Drosophila* embryo

**DOI:** 10.1101/079814

**Authors:** Maira Arruda Cardoso, Marcio Fontenele, Bomyi Lim, Paulo Mascarello Bisch, Stanislav Shvartsman, Helena Marcolla Araujo

**Affiliations:** Instituto de Ciências Biomédicas, Federal University of Rio de Janeiro; Institute of Molecular Entomology; Lewis-Sigler Institute for Integrative Genomics, Princeton University, Princeton, NJ; Instituto de Biofísica Carlos Chagas Filho, Federal University of Rio de Janeiro

**Keywords:** FκB, IκB, Toll, Cactus, Dorsal-Ventral patterning, Drosophila

## Abstract

The evolutionarily conserved Toll signaling pathway controls innate immunity across phyla and embryonic patterning in insects. In the Drosophila embryo Toll is required to establish gene expression domains along the dorsal-ventral axis. Pathway activation induces degradation of the IκB inhibitor Cactus resulting in a nuclear gradient of the NFκB effector Dorsal. Here we investigate how *cactus* modulates Toll signals through its effects on the Dorsal gradient and Dorsal target genes. Quantitative analysis using a series of loss and gain-of-function conditions shows that the ventral and lateral aspects of the Dorsal gradient behave differently respective to Cactus fluctuations. Unexpectedly, Cactus favors Dorsal nuclear localization required as response to high Toll signals at the ventral side of the embryo. Furthermore, N-terminal deleted Cactus mimics these effects, indicating that the ability of Cactus to favor Toll stems from mobilization of a free Cactus pool induced by the Calpain A protease. These results indicate that unexplored mechanisms are at play to ensure a correct response to high Toll signals.

**Summary:** The IκB protein Cactus favors high Toll signals, revealing that the ventral and lateral aspects of the Dorsal/NFκB nuclear gradient behave differently respective to Cactus concentrations in the Drosophila embryo.

## Introduction

The evolutionarily conserved Toll receptor pathway is implicated in the control of development, proliferation, and immunity. Toll signals are modulated at many levels, characteristic that warrants an assortment of possible outcomes (Mitchell et al., 2016). To understand Toll pathway architecture it is necessary to quantitatively define how each pathway element contributes to the final response. Inhibitor of NFκB (IκB) proteins comprise the Toll responsive complex together with NFκB effectors. Therefore, they are central elements of the Toll pathway that require careful analysis of their effects.

*Drosophila* is a unique system to identify how IκB proteins tune Toll responses as disturbances in IκB function can be analyzed concomitantly across a range of Toll activation levels. During embryogenesis, ventral-lateral activation of maternal Toll receptors leads to a ventral-to-dorsal nuclear gradient of the NFκB/c-Rel protein Dorsal (Dl). Inside the nucleus, differential affinity of Dl-target genes results in spatial control of gene expression along the dorsal-ventral (DV) axis (Rushlow and Shvartsman, 2012). High Toll signals lead to ventral mesodermal gene expression, intermediate signals induce lateral neuroectodermal genes, while low signals allow dorsal ectodermal gene expression. Toll signals are transduced through mobilization of adaptor proteins, protein kinase activation, and ultimately phosphorylation and proteasomal degradation of the sole *Drosophila* IκB protein Cactus. Cactus (Cact) degradation then exposes nuclear localization sequences in Dl and nuclear translocation ensues (Stein and Stevens, 2014).

The amount of nuclear Dorsal (nDl) is used as readout for the level of Toll pathway activation. Quantitative analysis of nDl in fixed and live embryos has shown that the Dl gradient is visible since early blastoderm cycle 9, increasing in amplitude with time until mitotic cycle 14 (Kanodia et al., 2009; Liberman et al., 2009; Reeves et al., 2012). Besides a deep understanding of the dynamics of the Dl gradient, how *cactus* affects this gradient has not been investigated. Using quantitative analysis we show that loss of *cactus* flattens the nDl gradient, implying that *cactus* favors Dorsal nuclear localization in addition to its widely established role to inhibit Toll signals.

## Results and Discussion

### Cactus favors Dorsal responses to high Toll signals

In order to identify how Cact tunes NFκB responses we investigated the effects of reducing *cactus* on Dl nuclear localization and expression of Dl target genes. It was previously reported that the nDl gradient expands, and the ventral *twist* (*twi*) expression domain widens in embryos generated from mothers carrying *cact* loss-of-function germline clones (Roth et al., 1991). This effect results from the near absence of Cact protein, which releases Dl inhibition in the cytoplasm. However, smaller reductions in *cact* lead to a range of DV embryonic phenotypes (Govind et al., 1993; Isoda and Nusslein-Volhard, 1994; Roth et al., 1991).

Here we investigate the effect of reducing, but not completely depriving the embryo of the Cact inhibitor. We progressively decreased maternal Cact protein to 50% wild type levels (Fig. 1 and Fig. S1). Reducing the *cact* gene dose by half leads to a minimal reduction in Cact protein and induces 30% embryo lethality (*cact*[A2] heterozygotes, (Govind et al., 1993), with no detectable DV embryonic phenotype (Fig. 1B,E,F,I,J and Table SI). Further Cact reductions, however, result in consistent alterations along the DV axis.

**Figure 1.**
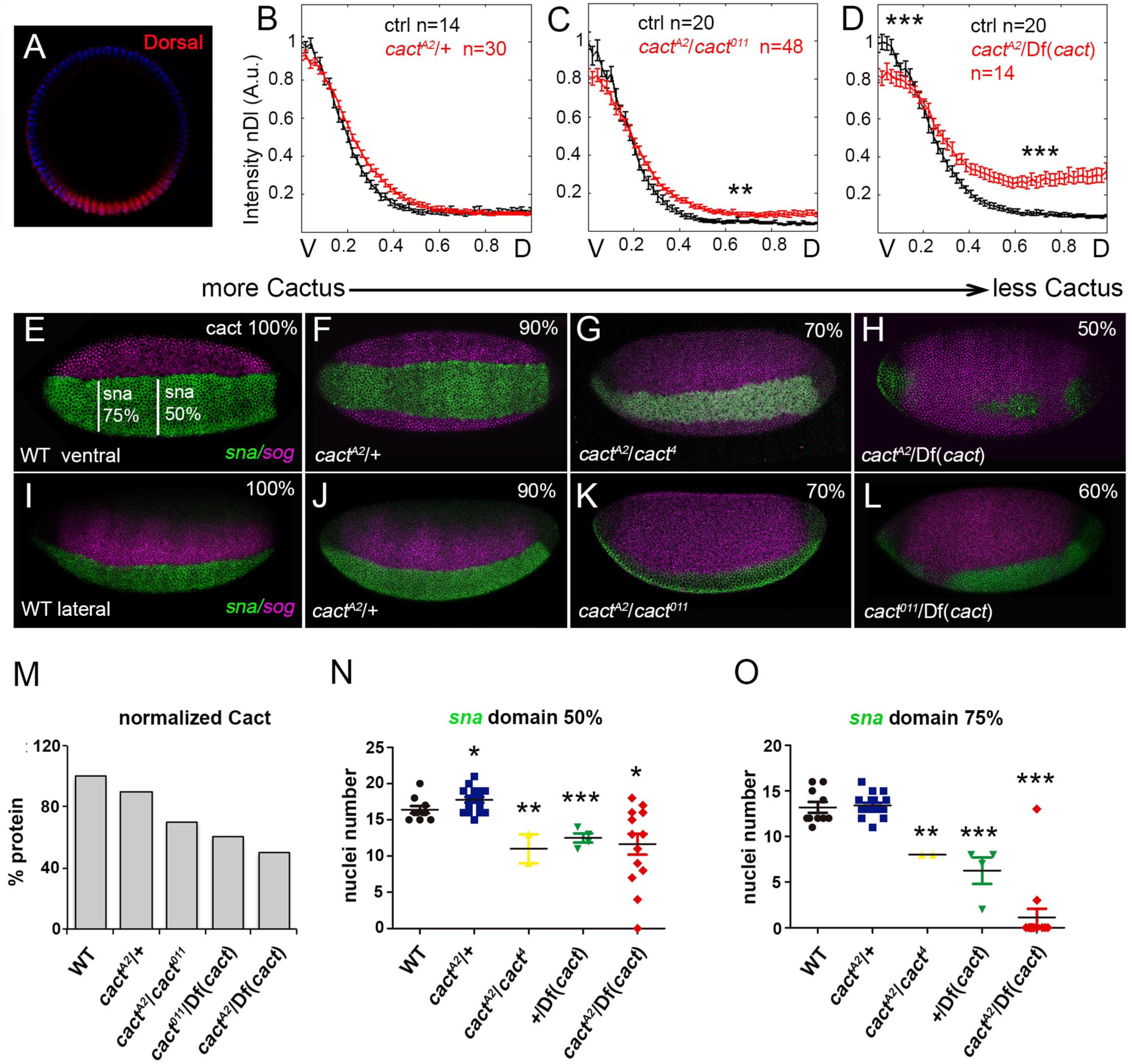
Cactus inhibits Toll in lateral regions and favors Toll signals in ventral regions of the embryo. A) Optical section of a control Dl gradient. B-D) Nuclear Dl levels were extracted and plotted as half gradients for control embryos (black) or embryos from *cact*[A2]/+ (B), *cact*[A2]/*cact*[011] (C) or *cact*[A2]/Df(*cact*) (D) mothers (red). Y axis represents fluorescence intensity of nDl along the ventral-to-dorsal embryonic axis (X axis). (E-L) *in situ* hybridization for *sna* (green) in the mesoderm and *sog* (pink) in the lateral neuroectoderm of embryos with maternal *cactus* reductions. In embryos from *cact* heterozygotes no change in the ventral and lateral territories are observed (F,J). Further reductions lead to a dorsal expansion of lateral *sog* and reduction in the ventral prospective mesoderm (G,H,K,L). Anterior is left, posterior right, dorsal up in G to L. Percentage in upper right indicates the average amount of Cactus protein relative to wild type, defined by quantification of protein levels in western blots (M and Suppl. Fig. 1). (N,O) Width of the *snail* domain measured at 75% and 50% egg length. Statistically significant differences based on Student's t-test, displayed as mean±s.e.m. (***P≤0.001, **P≤0.01, *P≤0.05).

In agreement with previous reports, decreasing Cact increases nDl in regions of the embryo that receive intermediate or low Toll signals (Fig. 1B-D): A 30% reduction in Cact protein is sufficient to expand lateral nDl levels to dorsal regions of the embryo (Fig 1C). A 40% reduction in Cact has the same effect, but additionally reduces nDl in the ventral domain (Fig. 1D). These changes in nDl are reflected on the pattern of Dl target genes. The lateral domain of *short gastrulation* (*sog*) expression, a gene that requires intermediate levels of nDl for activation (Stathopoulos and Levine, 2002), expands as Cact levels decrease by 30% or more (Fig. 1G,H,K,L). In the ventral prospective mesoderm the domains of *snail* (*sna*) and *twi* expression, genes that require high levels of nDl for activation (Ip et al., 1991; Papatsenko and Levine, 2005), are reduced as Cact levels drop (Fig. 1G,H,L and N,O and Fig. S2). Therefore, we find that Cact performs an additional function to favor Toll signals, distinct from its established function to inhibit Dl nuclear translocation. Importantly, this positive effect is restricted to regions that withstand high levels of Toll signaling.

Next, we investigated whether this positive effect depends on Dl. First, we reasoned that the amount of Dl protein might be limiting in contexts where Toll signals are high. In fact, in embryos laid by *dl-* heterozygous mothers nDl and the ventral *sna* domain narrow compared to controls (Fig. 2A,C) (Fontenele et al., 2013; Kanodia et al., 2009). Therefore, decreasing *dl* should sensitize the embryo to Cact reductions if the positive role of Cact depends on Dl. In agreement with this hypothesis, a minimal reduction in Cact (as in *cact*[A2] heterozygotes) further decreases nDl along the ventral and lateral domains of the embryo (Fig. 2B,D), an effect not observed in a wild type background (Fig. 1B). As a consequence of this reduction in Cact, the lateral *sog* domain expands into the ventral, high Toll activity, domain (Fig. 2C-H). These results confirm that *cact* favors Toll signals, and show that this effect is dependent on the amount of Dl.

**Figure 2.**
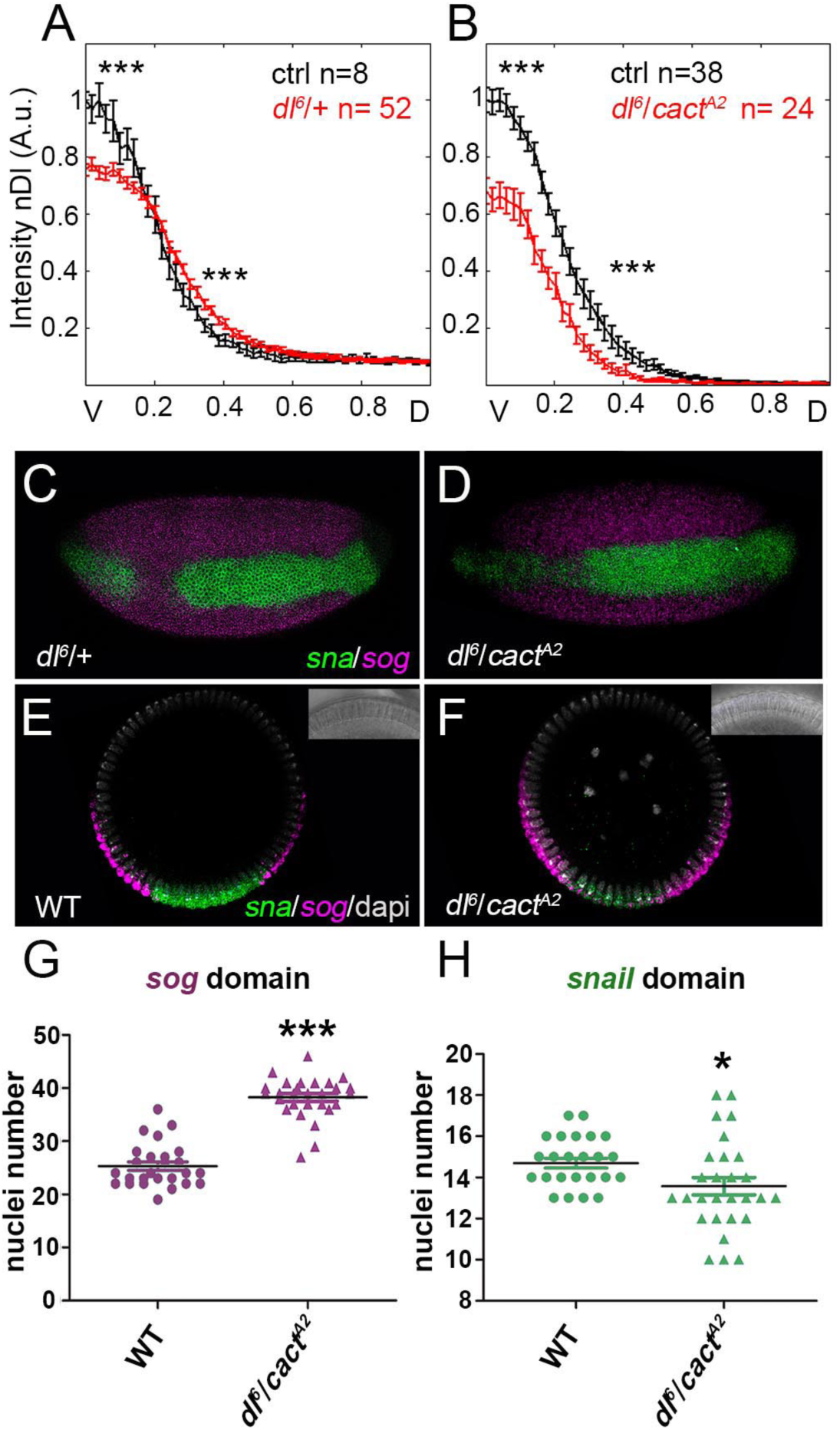
Cactus ability to favor Toll signals depends on the amount of Dorsal. A,B) nDl gradients for control embryos (black) or embryos from *dl*[6]/+ (A), or *dl*[6]/*cact*[A2] (B) mothers (red). The nDl gradient decreases by reducing maternal *dl* and *cact,* compared to control embryos stained concomitantly. (C-F) *in situ* hybridization and (G,H) quantification of *sna* (green) and *sog* (pink) in control (E) and maternal *dl*[6]/*cact*[A2] (F) transverse sections. Coincident *sog* and *sna* expression is seen in ventral regions (F). The size of the *sog* domain includes both left and right sides of the embryo since in *dl*[6]/*cact*[A2] this domain is frequently continuous. Statistically significant differences based on Student's t-test, displayed as mean±s.e.m. (***P≤0.001, *P≤0.05).

### Mobilization of free Cactus promotes Toll signals

Since decreasing *cact* lead to loss of ventral nDl, we asked whether we could detect a corresponding activity by overexpressing Cact constructs under the control of a maternal promoter (CaM,(Fernandez et al., 2001). Importantly, CaM>*cact-eGFP* perfectly recovers the nDl gradient in a *cact*[A2]/*cact*[011] loss-of-function background (Fig. S3A, compare to Fig. 1D), implying that maternal induction of *cact-eGFP* mimics wild type Cact behavior.

We tested the effects of two Cact constructs on the nDl gradient: full-length and N-terminal truncated Cact-eGFP. Two pathways control Cact levels: Cact is phosphorylated at N-terminal serine residues and degraded through the proteasome in response to Toll pathway activation (Bergmann et al., 1996; Fernandez et al., 2001; Hecht and Anderson, 1993; Reach et al., 1996; Shelton and Wasserman, 1993). Cact is also subject to C-terminal CKII kinase phosphorylation through a Toll independent pathway (Belvin et al., 1995; Liu et al., 1997; Packman et al., 1997). N-terminal truncated Cact mutants (*cact*[E10] and *cact*[BQ]) are irresponsive to Toll (Bergmann et al., 1996; Roth, 2001). Interestingly, N-terminal truncated Cact, hereafter referred to as Cact[E10], is produced endogenously from full-length Cact as a Calpain A protease cleavage product. Both full-length Cact and Cact[E10] bind Dl (Fontenele et al., 2013).

Maternal expression of Cact-eGFP and Cact[E10]-eGFP in *cact* loss-of-function backgrounds reveals that Cact[E10] favors Toll. When endogenous Cact is 30% reduced, Cact-eGFP recovers nDl and reduces lateral *sog* to a wild type pattern (Fig. S3A,F,I). Therefore, Cact-eGFP inhibits the expression of a Toll pathway target, as expected. The same level of Cact-eGFP expression has no effect once endogenous Cact is 50% reduced (Fig. 3A,C,D,F and G,HF,). Strikingly, Cact[E10]-eGFP increases the ventral *sna* domain and decreases the lateral *sog* domain in the same condition (Fig. 3A,B,D,E and G,H), although it has no effect when endogenous Cact is only 30% reduced (Fig. S3). Therefore, Cact[E10]-eGFP favors Toll in the ventral, high-Toll signaling domain and inhibits intermediate Toll signals in the lateral domain. In addition, this effect depends on the relative amount of endogenous full-length Cact.

**Figure 3.**
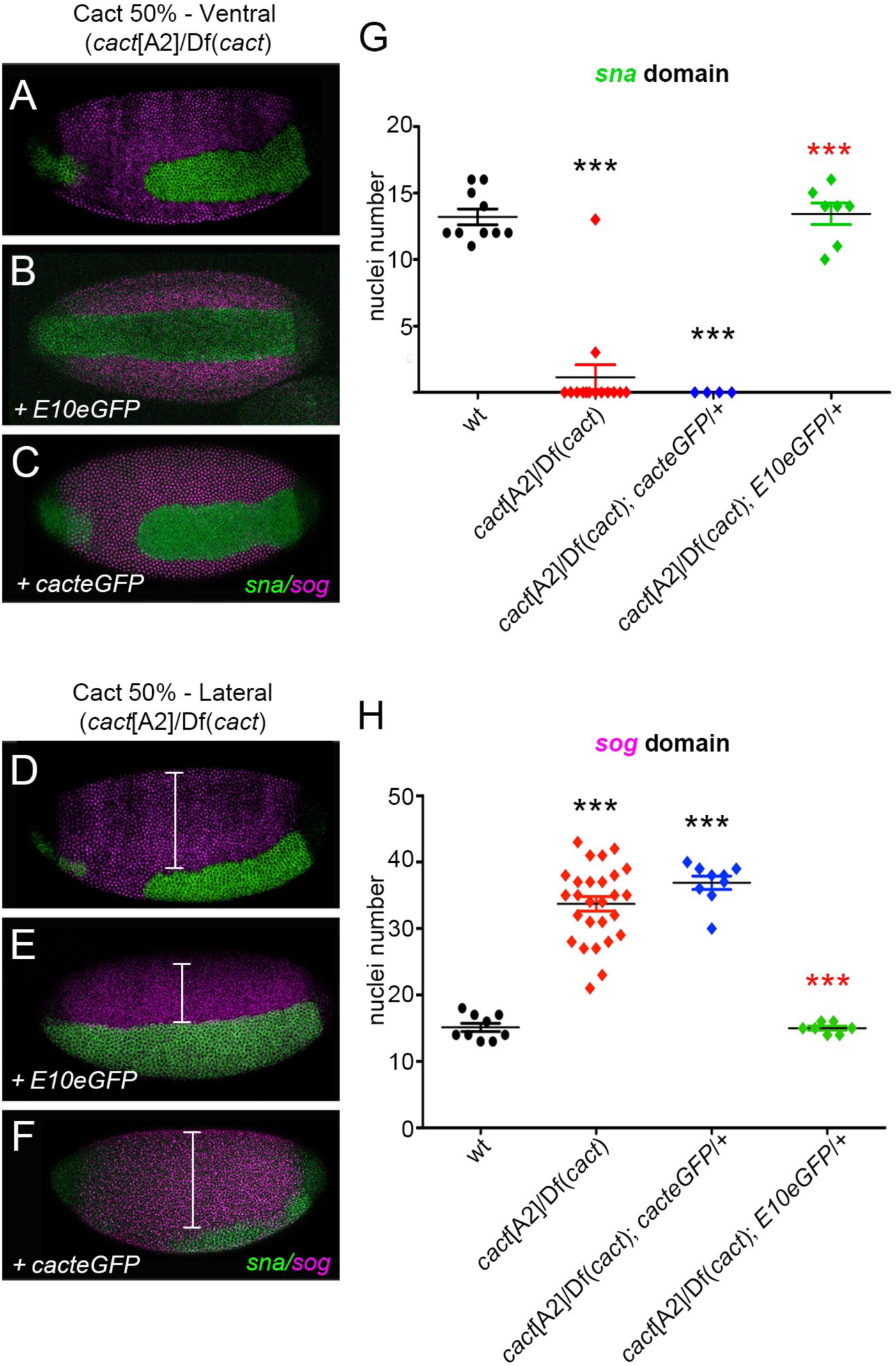
The Cact[E10] fragment favors Toll. (A-F) *in situ* hybridization for *sna* and *sog* in *cact*[A2]/Df(*cact*) (A,D); *cact*[A2]/Df(*cact*); *cactE10-eGFP*/+ (B,E) and *cact*[A2]/Df(*cact*); *cact-eGFP*/+ (C,F). (G,H) Quantification of *sna* at 75% egg length (G) and *sog* (H) domains. Full dorsal *sog* expansion is defined when the domain is greater that 25 cells and no dorsal *zen* expression is observed. Statistically significant differences based on Student's t-test, displayed as mean±s.e.m. (***P≤0.001).

The finding that Cact[E10] favors Toll is surprising considering that the gain-of-function *cact*[E10] allele inhibits Toll (Bergmann et al., 1996; Govind et al., 1993). Our results confirm that Cact[E10] inhibits Toll since decreases the lateral *sog* domain, target of intermediate Toll. However, Cact[E10] additionally favors Toll, depending on the respective amount of full-length Cact. Since N-terminal truncated Cact is produced endogenously (Fontenele et al., 2013), we conclude that Cact[E10] harbors the positive effect we have detected in the loss-of-function assays shown above.

The ability of Cact[E10]-eGFP to alter Toll signals depends on the amount of Dl. Cact-eGFP and Cact[E10]-eGFP produce a comparable decrease in nDl in a *dl-* heterozygous background (Fig. S4 compare to Fig. 2A), although Cact[E10]-eGFP has no effect in the presence of wild type Dl levels. Together with the results described above, this suggests that Cact[E10] competes with wild type Cact to bind Dl, and indicates the existence of a delicate balance between levels of full-length Cact, truncated Cact, and Dl for proper Toll signaling events.

In *Drosophila*, Cact partitions between free and NFκB-bound complexes. Dl-bound Cact comprises the Toll responsive complex (1Cact:2Dl), while Dl-free Cact (2Cact) is target of a Toll independent pathway (Bergmann et al., 1996; Liu et al., 1997). We have shown that the Calpain A protease, which leads to the production of Cact[E10], targets only Dl-free Cact. *Calpain A* knockdown decreases nDl in ventral regions, which implies that Calpain A favors Toll signals (Fontenele et al., 2013). Based on these findings, our results point to a model where Cact that is unbound to Dl (free Cact) is released by the action of Calpain A to favor Toll signals in ventral regions of the *Drosophila* embryo (Fig. 4). Interestingly, vertebrate Calpains also target IκB proteins and the pathways that control free and NFκB-bound Cactus and IκBαare conserved (Han et al., 1999; Li et al., 2010; Pando and Verma, 2000; Schaecher et al., 2004; Shen et al., 2001; Shumway et al., 1999). Therefore, our findings of a positive function for Cactus may have important implications for the control of vertebrate Toll signals.

**Figure 4.**
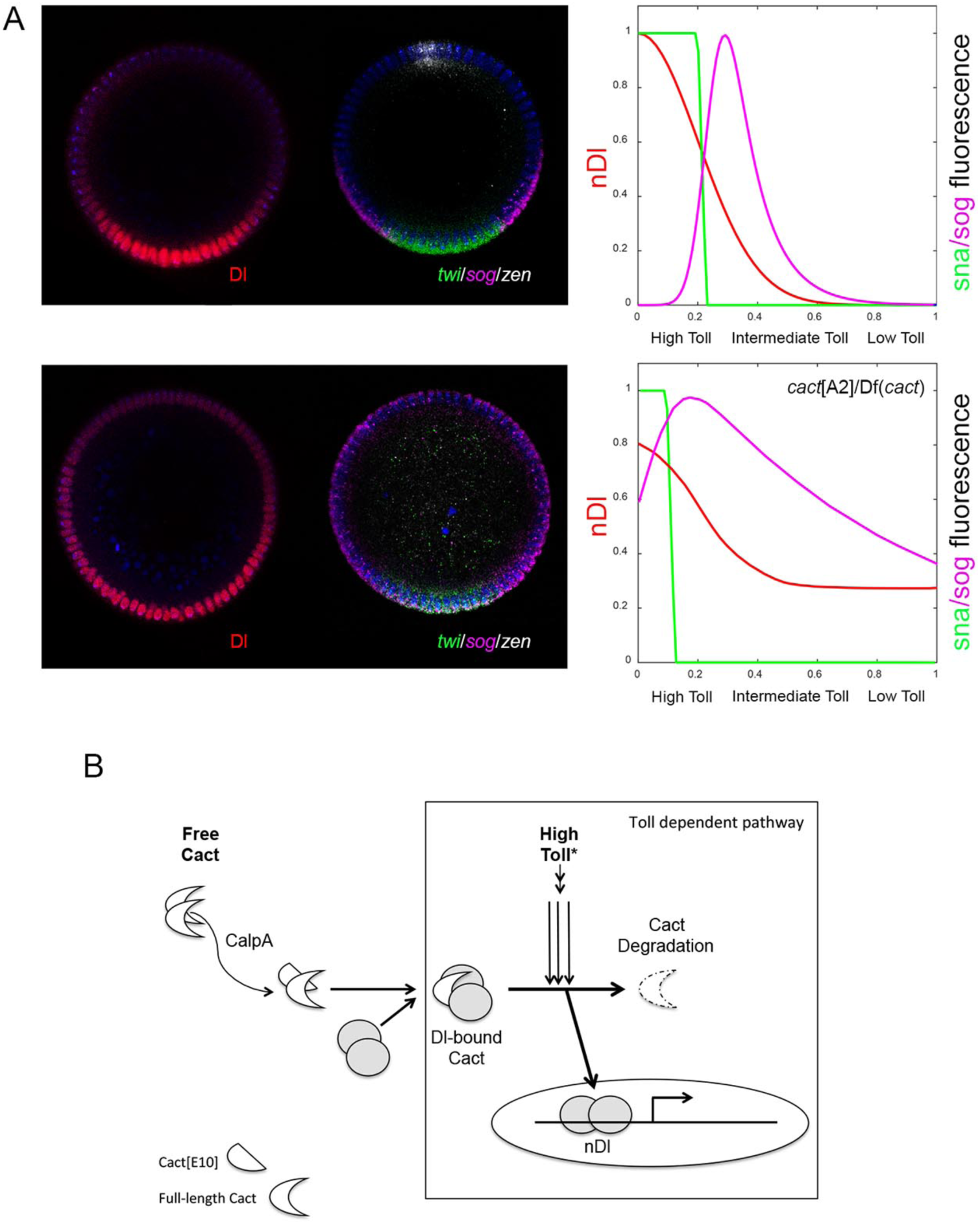
Model for free Cact function. A. nDl and ventral gene domains are reduced when Cact is decreased. B. This suggests that free Cact favors the formation of Dl/Cact complexes required for high Toll signals.

### On the nature of the positive function exerted by Cactus

Mathematical modeling pointed out the importance of IκB proteins in modulating Toll signals in vertebrates and in *Drosophila sp*. (Ambrosi et al., 2014; Kearns and Hoffmann, 2009; O’Dea et al., 2007). Particularly, it was shown that free IκB is an important regulatory target for the control of Toll signals (Konrath et al., 2014). Our previous analyses suggested that *Drosophila* Calpain A mobilizes free Cact to replenish limiting amounts of Cact:2Dl complexes for Toll signaling. With the present finding that Cact favors Toll, and that Cact[E10], a product of Calpain A activity, harbors this function, we now have quantitative data to support this hypothesis. Different mechanisms involving Cact[E10] could potentially favor Toll. For instance, Toll signals depend on the mobilization of pre-signaling complexes (Marek and Kagan, 2012; Sun et al., 2004), endocytosis (Huang et al., 2010; Lund et al., 2010), and how long NFκB proteins remain bound to DNA (Mitchell et al., 2016; O’Connell and Reeves, 2015). Thus, mechanisms involving Cact[E10] that favor the formation of pre-signaling complexes, the mobilization of endocytic vesicles containing Toll pathway elements or that alter the resident time of Dl in the nucleus, may positively impact Toll signals and explain our results. Furthermore, it has been recently proposed that Cact may function by a shuttling mechanism to concentrate Dl to the ventral side of the *Drosophila* embryo (Carrell et al., 2016) preprint), akin to the shuttling mechanism exerted by the BMP inhibitor Sog to shuttle and concentrate BMPs to the dorsal side of the embryo (Mizutani et al., 2005; Shimmi et al., 2005; Umulis et al., 2006). While further research is required to understand how Cact enhances nDl levels in ventral regions of the embryo and consequently Dl-target gene expression, we have clearly uncovered a novel function for the Cact inhibitor to favor Toll.

## Methods

### Fly stocks and genetic crosses

Lines used in this study were: loss-of-function *cact*[A2] and *cact*[011], generously provided by Steve Wasserman, *dl*[6] and *cact*[A4] obtained from the Bloomington Indiana Stock Center. Df(2L)*cact*[255]/CyO was used as a *cact* deficiency and is here denoted as Df(*cact*). Maternal overexpression lines were CaM>*cactus-eGFP* and CaM>*cactus*[E10]*-eGFP* as described in (Fontenele et al., 2013). All embryos were collected from mothers of the respective genotypes crossed to wild type Canton S males.

### Immunoblotting

Bleach dechorionated 30min-1h30min old embryos of the appropriate genotypes were homogenized in lysis buffer (1 embryo/l) and prepared for SDS-PAGE as in (Fontenele et al., 2009). Endogenous Cactus and Dorsal were detected with monoclonal antibodies from Developmental Studies Hybridoma Bank (DSHB, anti-Dl 1:100 and anti-Cact 1:500). Anti-αTubulin was used as loading control (DM1α; 1:3000; Sigma). Full length and truncated Cact-eGFP were detected using an anti-GFP polyclonal antiserum (1:1000; Novus Biologicals).

### Immunohistochemistry an in situ hybridization

For visualization of the nDl gradient mutant and control Histone-GFP embryos were mixed, fixed and processed concomitantly as in (Fontenele et al., 2013). Primary antisera used were monoclonal anti-Dl (7A4; 1:100; DSHB) and anti-GFP (1:1000; Novus Biologicals) to detect control gradients. Dl target genes were visualized by in situ hybridization as in (Fontenele et al., 2009).

### Quantitative analysis

Images of the nDl gradient and Dl target genes were collected from mid stage 14 embryos, as defined by the amount of membrane invagination around nuclei. Quantification of the Dl gradient was as described in (Kanodia et al., 2009) using Matlab, collected at 85% egg length using a microfluidic device to orient embryos for ends-on imaging (Chung et al., 2011). For upright imaging, a Nikon 60× Plan-Apo oil objective was used, and images were collected at the focal plane ∼90μm from the anterior pole of the embryo. For the overall effect on Dl target genes, embryos were imaged laterally. All genotypes were processed and analyzed in parallel, thus the same wild type control is shown in graphs. Images were acquired with a Nikon A1 or a Leica LSM confocal microscope.

### Statistical analysis

Student's *t*-test was performed for all experiments. Results are displayed as mean **± SEM**. The level of significance is shown in each figure (***P≤0.001, **P≤0.01, **P≤0.05),

## Acknowledgements

We thank Eric Wieschaus for the anti-twist antiserum. Monoclonal antibodies anti-Dl and anti-Cact originally developed by Ruth Steward were obtained from the Developmental Studies Hybridoma Bank, created by the NICHD of the NIH and maintained at The University of Iowa, Department of Biology, Iowa City, IA 52242.

## Competing interests

The authors declare that they have no conflict of interest.

## Author contributions

MAC, and MF performed experiments. MAC, MF and BO analyzed the data. MAC, PMB, SS and HMA developed the approach and prepared or edited the manuscript.

## Funding

This research was funded by the Brazilian Cientific Council (CNPq-Brazil, 477157/2013-0). MAC was a recipient of CAPES/Brazil local and overseas fellowships.

